# Insufficiency in airway interferon activation defines clinical severity to infant RSV infection

**DOI:** 10.1101/641795

**Authors:** Chin-Yi Chu, Xing Qiu, Matthew N. McCall, Lu Wang, Anthony Corbett, Jeanne Holden-Wiltse, Christopher Slaunwhite, Qian Wang, Christopher Anderson, Alex Grier, Steven R. Gill, Gloria S. Pryhuber, Ann R. Falsey, David J. Topham, Mary T. Caserta, Edward E. Walsh, Thomas J Mariani

## Abstract

Respiratory syncytial virus (RSV) is the leading cause of severe respiratory disease in infants. Other than age at the time of infection, the causes and correlates of severe illness in infants lacking known risk factors are poorly defined. We recruited a cohort of confirmed RSV-infected infants and simultaneously assayed the presence of resident airway microbiota and the molecular status of their airways using a novel method. Rigorous statistical analyses identified a molecular airway gene expression signature of severe illness dominated by excessive chemokine expression. Global 16S rRNA sequencing confirmed an association between *H. influenzae* and clinical severity. Interestingly, adjusting for *H. influenzae* in our gene expression analysis revealed an association between severity and airway lymphocyte accumulation. Exploring the relationship between airway gene expression and the time of onset of clinical symptoms revealed a robust, acute activation of interferon (IFN) signaling, which was absent in subjects with severe illness. Finally, we explored the relationship between IFN activity, airway gene expression and productive RSV infection using a novel *in vitro* model of *bona fide* pediatric human airway epithelial cells. Interestingly, blocking IFN signaling, but not IFN ligand production, in these cells leads to increased viral infection. Our data reveal that acute airway interferon responses are physiologically relevant in the context of infant RSV infection and may be a target for therapeutic intervention. Additionally, the airway gene expression signature we define may be useful as a biomarker for efficacy of intervention responses.

## Introduction

Respiratory Syncytial Virus (RSV), a negative strand RNA virus in the *Pneumoviridae* family, is the most important cause of respiratory tract infection during infancy, causing annual winter outbreaks in the US^1–6^. In the US approximately half of the 4 million newborns are infected during their first winter with 1-3% hospitalized and an additional 4-7% and 10-16% seen in emergency departments or physician offices, respectively, for RSV infections^7^. Mortality is uncommon in the US (~50 deaths annually); however, in developing countries it is estimated that annually RSV causes 118 thousand deaths, 6 million cases of severe acute lower respiratory illness and 3 million hospitalizations in children under 5 years of age^8,9^. Currently there is no available vaccine for RSV in infants, although several candidate vaccines are in clinical trials.

Long established major risk factors for severe illness include prematurity, bronchopulmonary dysplasia, cyanotic heart disease, neuromuscular disease, and immune compromise^4,10^. However, approximately 70% of hospitalized infants in the US have no overt risk factors for severe illness. Although young age at infection, environmental influences such as tobacco smoke exposure, viral load and strain, low levels of maternally derived RSV-neutralizing antibody, as well as a multitude of genetic host factors have been associated with severe disease in some but not all studies^4, 11–22^. Recently, the presence of *Haemophilus influenzae and Streptococcus pneumoniae* in the nasal microbiota during RSV infection has been associated with greater severity^23–25^. Finally, and importantly, the infant’s immune response to RSV is thought to be a major driver of disease pathogenesis, especially during primary infection^26,27^. Several studies in infants suggest that Th2 biased responses and Th17 responses during primary infection may contribute to a more inflammatory and severe outcome^28–30^. Innate immune responses by immune cells, such as neutrophils, and respiratory epithelial cells are also likely to play pivotal roles in both eliminating virus replication as well as enhancing or moderating the inflammatory response^31,32^.

The AsPIRES (<u>As</u>sessing and <u>P</u>redicting Infant <u>R</u>SV <u>E</u>ffects and <u>S</u>everity) study is a comprehensive study designed to identify factors associated with disease severity in full term healthy infants less than 10 months of age undergoing primary RSV infection. In this report we analyze gene expression of nasal respiratory epithelial cells, in addition to the influence of respiratory microbiota, in relation to illness severity during primary RSV infection. In addition, we describe the development of an *in vitro* model of RSV infection to explore the relationship between interferon activity, airway cell gene expression and productive RSV infection.

## Methods

### Study subjects

RSV infected infants were identified and enrolled into the Assessing Predictors of Infant RSV Effects and Severity (AsPIRES) study during three winter (2013-14, 2014-15, and 2015-16) in Rochester NY as previously described^33,34^. All infants were less than 10 months of age at the time of primary RSV infection and came from one of three cohorts; (1) infants prospectively enrolled at birth and followed for development of RSV infection, infants enrolled upon hospitalization with RSV infection, and (3) infants identified with RSV in emergency rooms or physician offices managed as outpatients. All subjects were previously healthy full-term infants (gestational age ≥ 36 weeks) born after May 1 of the previous spring. Infants hospitalized only for apnea were excluded as well as those with any known high-risk conditions. The Research Subject Review Board of the University of Rochester and Rochester General Hospital approved the study and all parents provided written informed consent.

### Study Protocol and Procedures

RSV infection was identified by reverse transcription polymerase chain reaction (RT-PCR) on nasal swabs. Following confirmation of RSV infection all study subjects underwent a standard evaluation at three time points. The first illness visit occurred within 24 hours of RSV diagnosis, a second visit 12-16 days after illness onset, and final visit on illness day 25-32. Demographic and clinical data were collected at each visit. At the first and third visits a nasal swab was collected from one nostril for analysis of respiratory microbiota, followed by collection of nasal epithelial cells from the contralateral nostril as previously described^35^. Briefly, after washing the nares with 5 ml of sterile saline to remove mucus and cellular debris the mid turbinate was brushed for 5 seconds with a flocked swab (Copan, FLOQSwabs™ catalog # 525CS01, Copan, Murrieta, CA) to remove respiratory epithelial cells. The swab was immediately placed in 2 ml of RNA protect and maintained at 4°C until the cells were recovered by filtration and homogenized in RNA lysis buffer, as described^35^.

### Statistical Analysis of Clinical Data

Descriptive characteristics of the study cohort were reported in Table 1. For binary variables, percentages and frequencies were reported; for continuous variables, means and standard deviations were reported. Appropriate statistical tests were performed to test the association between each clinical variable and disease severity. Specifically, for dichotomous severity (mild v. severe), we used two-sample Welch t-test and Fisher’s exact test for continuous and binary variables, respectively; for continuous severity (GRSS), two-sample Welch t-test and Pearson correlation test were used for the binary and continuous clinical variables, respectively. A p-value <0.05 was considered statistically significant. All analyses were performed with SAS (version 9.3; SAS Institute, Cary, NC) and the R programming language (version 3.2; R Foundation for Statistical Computing, Vienna, Austria).

**Table 1.** Subject Demographics. Data represent samples collected at the time of acute infection (1-10d after onset of clinical symptoms) from 106 subjects. Reported in this table are (mean±STD) for continuous variables, and the frequency in both groups for categorical variables. P-values reported in this table were computed by statistical tests that are appropriate to the nature of the variables (see Section “Statistical Analysis of Clinical Data” for more details). Only subject age at time of infection was significantly associated with clinical severity, and only when severity is defined on a continuous scale.

| VARIABLE | n=106 | P-value<br>Mild (42) vs. Severe (64) | P-value<br>GRSS (4.35±2.66) |
| --- | --- | --- | --- |
| Age, months | 3.35±2.22 | 0.5122 | **0.0323** |
| Gestational Age, weeks | 38.90±1.37 | 0.3437 | 0.3064 |
| Birth Weight, kg | 3.34±0.61 | 0.7468 | 0.4625 |
| Family size | 4.16±2.25 | 0.3703 | 0.6150 |
| Gender, Female / Male | 54 (51%) / 52 (49%) | 0.4275 | 0.5840 |
| Caucasian (only), True / False | 62 (58%) / 40 (38%) | 0.1495 | 0.5832 |
| Tobacco smoke exposure, True / False | 36 (34%) / 70 (66%) | 1 | 0.4492 |
| RSV Strain, A / B | 59 (56%) / 46 (43%) | 0.6514 | 0.8365 |
| Viral co-infection or bacteria colonization, True / False | 83 (78%) / 23 (22%) | 1 | 0.3190 |
| Bacterial colonization, True / False | 79 (75%) / 27 (25%) | 0.6498 | 0.2092 |

<sup>a</sup>; all values are mean ± SE unless otherwise noted
<sup>b</sup>; severe disease as defined by hospitalization status

### Defining Illness Severity

The severity of the clinical illness in these patients was defined by a global respiratory severity score (GRSS)^33^. Briefly, a Global Respiratory Severity Score (GRSS) was defined by a weighted combination of 9 clinical measures including general appearance, presence of rales, wheezing, cyanosis, retractions, lethargy, poor air movement, maximal age-adjusted respiratory rate, and worst room air oxygen saturation. The GRSS is defined on a continuous scale from 0-10, and is highly correlated with need for hospitalization. For some secondary analyses, severe illness was dichotomously defined as GRSS>3.5.

### Library Preparation and Sequencing

The RNA was isolated and assessed for quality and quantity (average yield 275 ng per sample). 1 ng of total RNA was amplified using the SMARter Ultra Low amplification kit (Clontech, Mountain, CA) with PCR amplification to generate sufficient cDNA for library construction. Amplified cDNA quantification was determined with the Qubit Flourometer (Life Technologies, Grand Island, NY) and quality was assessed using the Agilent Bioanalyzer 2100 (Santa Clara, CA). Library construction was performed using NexteraXT library kit (Illumina, San Diego, CA) with 1 ng of input amplified cDNA per manufacturer’s recommendations. Nextera libraries were quantified with the Qubit Flourometer (Life Technologies, Grand Island, NY) and quality was assessed using the Agilent Tape Station (Santa Clara, CA). Libraries were sequenced on the Illumina HiSeq2500 (Illumina, San Diego, CA) to generate 20 million 1X100-bp single end reads per sample^35^.

### Read Mapping, Data Normalization and Filtering

Sequences were aligned by STAR against human genome version of GRCh38 and normalized to FPKM. We used the following non-specific filtering strategy to remove genes with low expression values indistinguishable from background signal. First, we compute *M*_*i*_, the 95% sample quantile of the expression level of the *i*th gene, for all genes. Next, we defined trimmed maximum log-expressions as *L*_*i*_ = log_2_(*M*_*i*_ + 1), and removed those genes with *L*_*i*_ < 0 (or equivalently, *M*_*i*_< 7.0). After this filtering step, a total of 13,819 genes were selected for further analyses. After removing fifteen samples with total number of mapped reads less than 5 million and ten samples with low average correlation with other samples, 175 samples were selected for further analyses. Objective metrics for data from these samples suggested a high degree of quality (Supplemental Figure 1), similar to what we have previously published^35^.

Due to the fact that our study spanned three years, all samples were processed as six library batches. We noticed that there are significant batch effects in the total number of mapped reads of these 175 samples. In addition, using an analysis of variance (ANOVA) *F*-test with false discovery rate (FDR) controlled at 0.05 level, we found 3,984 genes (28.8% of the expressed transcriptome) had significantly different mean expression across batches. Based on these observations, we used ComBat^36^ to remove batch effects. As expected, after applying ComBat, no gene had a significant batch effect based on an ANOVA *F*-test. To avoid spurious findings due to outliers, we also winsorized the data at 1% and 99% levels. Specifically, if an observation was less (greater) than 99% of the data, we replaced its value by the 1% (99%) sample quantile. We also removed 131 genes for which their expression was winsorized in more than half of the samples (n>53). In summary, a total of 13,688 gene expression profiles passed the quality control procedure and were selected for further investigations.

### Gene Significance Analyses

Both univariate and multivariate gene significance analyses were conducted by using the R package LIMMA^37^ with a robust M-estimator^38^. Nine covariates (global respiratory severity score (GRSS), sex, race, gestational age, visit age, time since disease onset (time), smoking exposure, virus co-infection, and bacterial colonization) were investigated via univariate analyses, in which the association between a single covariate and gene expression was estimated. Following the univariate analyses, we conducted a multivariate analysis that included the same nine significant clinical variables. In addition, we also tried a simplified multivariate regression model with the five most significant covariates (time, GRSS, sex, race, and bacterial colonization) identified in the full multivariate regression analyses. In both cases, regression t-tests were used to determine the significance of linear associations between covariates and gene expression levels. The Benjamini-Hochberg multiple testing correction^39^ was applied to control the false discovery rate (FDR) at 0.05^39^.

### Detection of Pathogenic Virus and Bacteria (TLDA)

Initial diagnosis of RSV infection was confirmed by RT-PCR as described^11^. TaqMan® Array Card (TAC) technology was used to detect other common respiratory pathogens in the initial nasal swab as well as *Hemophilus influenzae and Streptococcus pneumoniae*, as previously described^40–42^. Viral targets included influenza A and B, respiratory syncytial virus (RSV), human metapneumovirus (hMPV), parainfluenza virus (PIV) 1, 2 and 3, human rhinovirus (hRV), human enterovirus (EV), human parechovirus (hPeV), coronavirus 1 through 4, (229, NL63, OC43, and HKU1 respectively), adenovirus (ADV), and human bocavirus (hBoV). *H. influenzae, S. pneumoniae, M. pneumoniae, C. pneumoniae, B pertussis, M. hominis, and U Ureaplasma* were also included on each card^35,40^. Detection of the human RNase P (RNP3) gene was a positive control to confirm the presence of cellular material.

### Microbiome Analysis

Airway microbiota analysis from infants was performed essentially as previously described^35^. Briefly, nasal specimens were collected from the anterior nares during the acute illness visit. V3-V4 16S rRNA was amplified from total genomic DNA extracted from the swab and sequenced (2 × 300bp) on an Illumina MiSeq (Illumina, San Diego, CA). 16S rRNA bacterial sequence reads were assessed for quality and analyzed using phylogenetic and Operational Taxonomic Unit (OTU) methods in the Quantitative Insights into Microbial Ecology (QIIME) software^43^, version 1.9. For the purposes of OTU relative abundance analysis, the raw OTU table was normalized using the cumulative sum stabilization method from the `metagenomicSeq` R package^44^.

### Transcriptome and Microbiome Integration Analyses

We limited analyses to those 83 subjects for which both transcriptome and microbiome samples were available, and who were not on antibiotics. Data were available for a total of 471 operational taxonomic units (OTUs). After filtering those detected in <3% of subjects, and those for which neither genus or species was discernable, 148 OTUs were used for analysis. A linear regression of GRSS as outcome was conducted to estimate the association between GRSS and each OTU, while controlling for visit age. The results showed two highly correlated OTUs, both for *Haemophilus taxa (*species *Influenzae* and species unknown). Further analyses combined these OTU as *H. Influenzae.* Similar regressions were performed for additional clinical variables. Next, we tested for a univariate association between GRSS and gene expression while adjusting for visit age. This identified 1185 genes associated with GRSS based on a regression t-test (unadjusted p value<0.05). Finally, a multivariate linear regression was conducted on GRSS as the outcome variable and including visit age, combined *H. Influenzae* and each of the 1185 selected gene expression profiles.

### Nasal Fluid Cytokine Determination Multiplex Assay

Nasal wash samples were tested for cytokines and chemokines using R&D Systems Human Luminex human assay (Minneapolis, MN) with magnetic beads. Preparation of cytokine standards and procedure of the assays were carried out following the manufacturer suggested protocol. The beads were read using MAGPIX (Luminex Corp., Austin, TX). xPONENT software (Luminex Corp., Austin, TX) was used for instrument control, data acquisition, and data analysis. The 14 cytokines and chemokines measured were IL-6, IL-16, IL-17, IFN-λ 2 & 3, IL-33, IFN-β, CCL3/MIP1α, CCL7/MCP3, CCL20/MIP3α, CXCL1/GROα, CXCL2/GROβ, MMP1 & TRAIL. Statistical analysis was carried out for non-paired samples with the Mann-Whitney U test.

### Functional Classification

Genes identified as differentially expressed were subsequently used for canonical pathway identification and upstream regulator analysis using Ingenuity Pathway Analysis (IPA; http://analysis.ingenuity.com/pa/); ontology and phenotype functional enrichment analysis using ToppGene Suite (https://toppgene.cchmc.org/); cell-type enrichment identification using CTen (http://www.influenza-x.org/~jshoemaker/cten/advanced_example.php).

### Gene Expression Validation

Quantitative real-time polymerase chain reaction (qPCR) was performed using aliquots of RNA samples obtained for sequencing. cDNA was generated from approximately 250 ng RNA using the iScript cDNA Synthesis Kit (BioRad, Hercules, CA) according to manufacturer’s recommendations. PCR was performed using gene-specific primer sets (http://pga.mgh.harvard.edu/primerbank/) and Taqman chemistry (Universal Master Mix II with UNG, Life Technologies) on a ViiA 7 Real-Time PCR System (Life Technologies). Gene expression levels were calculated relative to PPIA (cyclophilin A) using the ddCT method.

### Development of *in vitro* RSV infection model using respiratory epithelial cells

Pediatric/Infant Human Lung Epithelial (PHLE) cells were generated from transplant donor quality lung from the LungMAP program as we recently described^45^. Briefly, fresh lung tissues were digested with a protease cocktail. Single cell suspensions were washed 2 times with DPBS supplemented with 1% Penicillin-Streptomycin (Gibco), 50 µg/ml Gentamicin and 0.25µg/ml amphotericin B and centrifuged at 800 xg for 10 minutes and grown in SAGM supplemented with 1% FBS. After 24 hours, non-adherent cells were removed. Epithelial cells were expanded and fibroblast-like cells were removed by treatment with 0.0125% trypsin (with EDTA) in DPBS at room temperature. For establishing Air-liquid interface cultures, 100,000 cells were seeded on rat tail collagen I (34.5µg/mL) coated Transwell inserts (12 well PET membrane, 0.4μm pore size, 12 mm diameter). After 24-48 hours, culture medium was changed to 1:1 mixture of BEGM/DMEM. When cells reached confluence, with appropriate resistance (≥ 300 Ohms*cm2), they were transferred to ALI by removing the apical medium. Upon transition to ALI, cells were maintained in PneumaCult-ALI medium containing supplements and hydrocortisone, according to manufacturer’s instructions. ALI cultures were differentiated for 10-14 days at ALI before virus infection.

RSV GFP-A2 strain (gift of Dr. Mark Peeples) was generated in Hep2 cells as previously described^46^. ALI-differentiated PHLE cultures were washed twice with PBS and infected apically with RSV at a multiplicity of infection of 1. After one hour at 37°C, the inoculum was removed and the apical surface was rinsed with PBS twice to remove unbound viral particles. Infected PHLE were maintained at ALI for up to 48 hours, and monitored for infection using a fluorescence microscope. After 48 hours, cells were harvested and RNA isolated, and subjected to qPCR as described above. Infection was quantified by measuring GFP fluorescence with ImageJ software (https://imagej.nih.gov/ij/) and RSV-M gene quantification using real-time polymerase chain reaction (qPCR). Where indicated, interferon ligands (30 ng/ml IL28 or 2.5 ng/ml IFN-β; R&D Systems) were provided to PHLE cells for an hour prior RSV infection and maintained throughout the infection experiment. Where indicated, interferon signaling inhibitors (4 µM Ruxolitinib or TPCA-1; Selleck Chemicals) were provided to PHLE cells for an hour prior RSV infection and maintained throughout the infection experiment.

## Results

### Subject Demographics

We sought to understand target organ resident cell responses during primary RSV infection in the first year of life in infants displaying the full spectrum of illness severity. We implemented our recently described nasal cell sampling procedure^35,47^ to measure infant airway transcriptional responses. Of the 139 RSV infected infants enrolled in the AsPIRES study, 106 had satisfactory nasal samples and gene expression results for analysis. Demographic data of these 106 infants are described in Table 1. Infants ranged from 0.5 to 9.4 months of age (mean age 3.3 months) and were equally distributed by gender. Sixty-three (59%) were hospitalized during the infection while the other 43 were managed as outpatients.

Subjects were assigned a continuous Global Respiratory Severity Score (GRSS) as previously described^33^. There was no association between severity (defined as being hospitalized or by GRSS) and RSV strain, family-reported environmental tobacco smoke exposure, or the presence of other viral pathogens or bacterial colonization based on a positive RT-PCR (Table 1).

### Airway Transcriptional Correlates of Disease Severity

Transcriptome analysis of airway samples collected from subjects during the acute phase of their RSV infection included the assessment of 13,688 genes following standard QC-based sample and gene filtering^28,35^. As we previously described, gene expression data from this procedure is enriched in canonical (e.g., CDH1, EPCAM) and upper airway-specific (e.g., BPIFs, MUCs) marker genes, with lower levels of expression for leukocyte markers (Supplemental Figure 2). We initially explored the relationship between gene expression and demographic variables. Univariate analysis indicated that many variables were not associated with substantial differences in gene expression after appropriate adjustments for multiple testing (Table 2). As previously reported^35^, the presence of known bacterial pathogens had a large impact on airway gene expression (n=470 genes), appearing to be driven more by the presence of *S. pneumoniae* (n=691 genes) than *Moraxella* or *H. influenzae* when identified by standard RT-PCR microbiologic diagnostic assays (TAC cards, as described above). Notably, the time elapsed from the onset of clinical symptoms (time) also had a major impact on gene expression (n=216 genes). Interestingly, the presence of an additional pathogen during acute infection, in the aggregate, was not significantly associated with clinical severity (P>0.319) (Table 1).

**Table 2.** Number of genes with significant changes associated with individual clinical and demographic variables. We performed univariate & two multivariate regression models to identify gene expression changes associated with intrinsic (age, GRSS, days since onset of illness) and extrinsic (tobacco smoke exposure and bacterial colonization or viral co-infection) factors. Shown are the number of genes identified as significant for each variable in the model.

|  | Clinical Variables | n=106 | Univariate<br>(BH p<0.05) | Full<br>multivariate<br>(BH p<0.05) | Simplified<br>multivariate<br>(BH p<0.05) |
| --- | --- | --- | --- | --- | --- |
| **Continuous**<br>**Variables** | **Clinical Severity (GRSS)** | 4.35±2.66 | 142 | 252 | 308 |
|  | **Days Since Onset of Illness, day** | 4.60±0.20 | 216 | 728 | 617 |
|  | **Gestational Age, weeks** | 38.90±1.37 | 0 | 0 |  |
|  | **Age, months** | 3.35±2.22 | 5 | 1 |  |
| **Categorical**<br>**Variables** | **GRSS , Mild/Severe** | 42 (40%)/64 (60%) | 0 |  |  |
|  | **Gender, Female/Male** | 54 (51%)/52 (49%) | 18 | 60 | 21 |
|  | **Caucasian, True/False** | 62 (58%)/40 (38%) | 13 | 17 | 19 |
|  | **Tobacco Smoke Exposure, True/False** | 36 (34%)/70 (66%) | 0 | 0 |  |
|  | **Viral co-infection, True/False** | 18 (17%)/88 (83%) | 7 | 3 |  |
|  | **Bacterial colonization, True/False** | 79 (75%)/27 (25%) | 470 | 135 | 300 |
|  | ***S. Pneumoniae*, True/False** | 39 (37%)/67 (63%) | 691 |  |  |
|  | ***H. Influenzae*, True/False** | 29 (27%)/77 (73%) | 22 |  |  |
|  | ***Moraxella*, True/False** | 67 (63%)/39 (37%) | 0 |  |  |

Gene expression displayed an interesting association with clinical severity. No gene was marginally identified as significantly associated with severity (after adjustment for multiple testing adjustment), when severity was defined dichotomously as mild and severe. When severity was defined continuously by GRSS, 142 genes were significantly associated with GRSS. In order to better define gene expression changes independently associated with GRSS, we completed a multivariate analysis including sex, race, bacterial colonization and time since onset of clinical symptoms in the model. This analysis identified 252 genes significantly associated with disease severity (Supplemental Figure 3). In this multivariate model, 728 genes were associated with time since illness onset and 135 genes were associated with the presence of any bacterial pathogen (Table 2).

Genes significantly associated with severity, in both the univariate (data not shown) and multivariate models were rich in chemokines, cytokines and interleukin-related molecules (Supplemental Figure 4). Ontological analysis of this gene set identified multiple biological functions related to viral infection (Fig. 1A), confirming the validity of the approach and the data. Interestingly, this analysis suggested that gene expression biomarkers of severity are associated with pneumonia (Fig. 1B). This observation, consistent with the lower respiratory track pathogenesis of severe RSV infections, also supports the utility of upper airway transcriptome as a surrogate for identifying relevant disease mechanisms in the lung. We also assessed cell signatures in these severity associated gene expression biomarkers. Unlike the epithelial predominant signature of the healthy asymptomatic infant^35^, severe clinical responses to RSV infection were associated with signatures of multiple blood and inflammatory cells, as well as smooth muscle (Fig. 1C). The most predominant cell signatures were those related to CD14+ and CD33+ myeloid lineages. Pathway analysis confirmed increases in expression of myeloid cell-related genes in severe cases, and also strongly implicated IL17-related responses (Fig. 1D).

**Figure 1.**
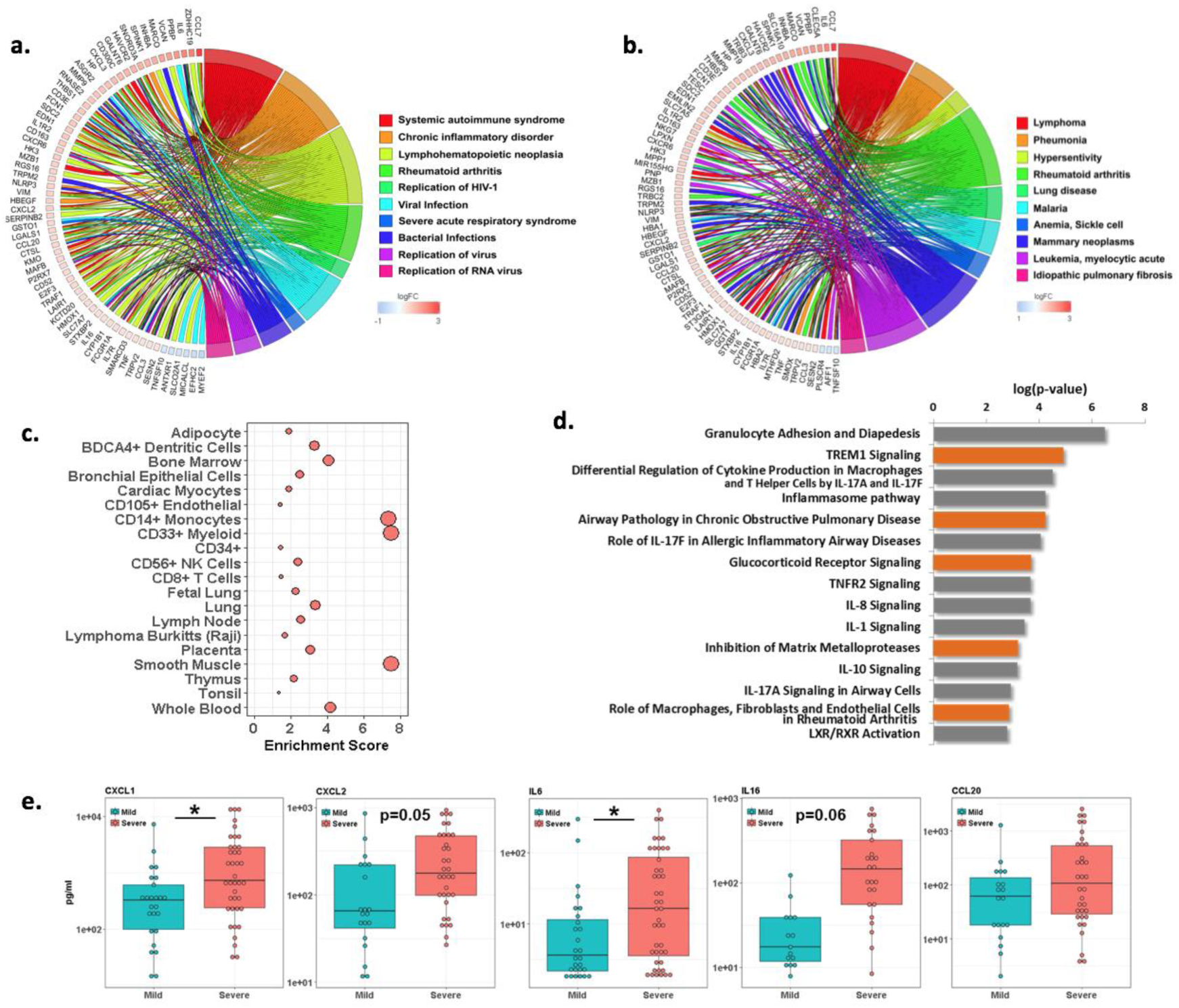
Airway gene expression patterns associated with clinical severity in RSV-infected infants. Gene expression from nasal/airway samples, collected from 106 infants infected with RSV, was assessed by high throughput RNA sequencing. Circos plots displaying biological function (a.) and disease (b.) ontologies associated with severity. Shown are a subset of genes with expression patterns significantly associated (FDR<0.05, fold change>1.5) with clinical severity, following adjustment for other variables in a multivariate model, and key ontologies significantly associated with severity (n=629 genes, FDR<0.05). Biological functions are enriched in viral infection-related ontologies, and disease ontologies include pneumonia, a lower airway pathology. Cell type enrichment plots (c.) indicate severity genes are associated with multiple hematopoietic lineages, predominantly CD14+ monocytes, CD33+ myeloid cells and smooth muscle cells. Canonical pathway analysis of severity-associated genes (d.) suggests alterations in immune signaling in infants with severe clinical symptoms including IL17. Multiplex ELISA analysis of nasal washes from RSV infected infants (e.) confirms that the production of select immune-related proteins are increased in infants with severe clinical symptoms.

We validated severity-associated gene expression changes at the transcriptional level by qPCR of RNA samples from the same 106 subjects (Supplemental Table 1). We also performed multi-plex ELISA analysis, using nasal washes obtained immediately prior to nasal brush sample collection, to confirm the biological impact of these transcriptional changes. As shown in Figure 1E, we validated significant differences in the level of multiple inflammatory biomarkers associated with severity (eg, CXCL1, CXCL2, IL6), and non-significant differences in others (eg, IL16).

### Effects of Airway Microbiota on Severity Biomarkers

Recent work by others has demonstrated the specific presence of *H. influenzae* in the airway as a correlate to severe clinical responses in infants with RSV. We completed unbiased microbiota analysis of the airway of our subjects, using 16S rRNA sequencing of nasal swab samples simultaneously obtained from contralateral nares. We calculated operational taxonomical units (OTUs) for all identifiable organisms, and individually assessed the relative abundance of 148 OTUs for association with clinical severity. *H. influenzae* (Supplemental Figure 5), was the only OTU significantly associated with severity following adjustment for multiple testing in this cohort (Supplemental Table 2). As can be seen in Figure 2A, although a great degree of variation in relative abundance of *H. influenzae* existed within both groups, more severely affected infants on average had a much higher abundance. Although undefinable using 16S rRNA data, presumably this represents non-typable *H. influenzae* (NTHI) since the incidence of *H. influenzae type B* is uncommon. Interestingly, 9 OTUs (not including *H. influenzae*) were significantly associated with age, suggesting the nasal microbial communities evolve extensively during infancy in this cohort^48^.

**Figure 2.**
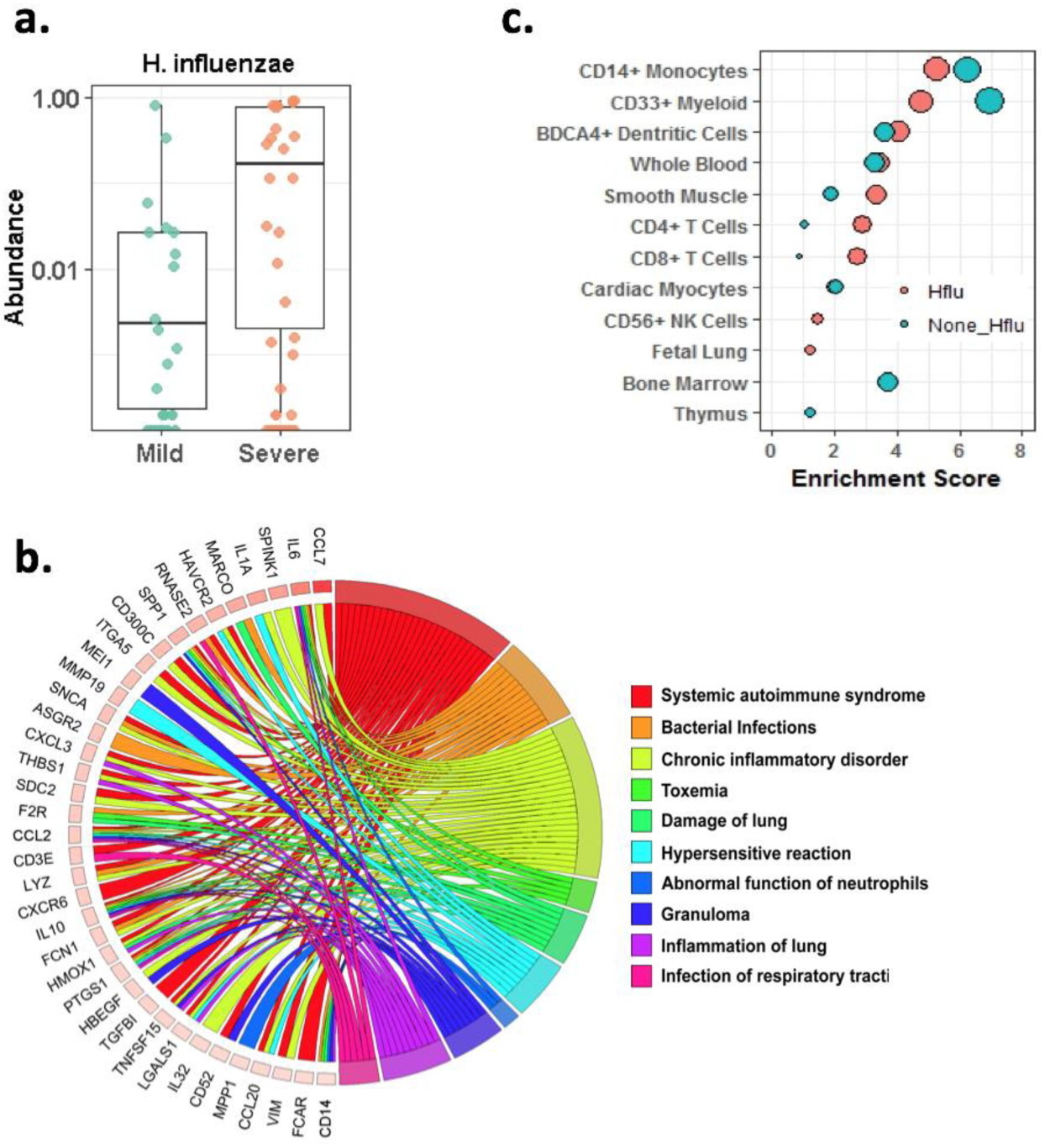
Microbiota effects on airway gene expression in RSV-infected infants. 16s rRNA sequencing was used to interrogate the airway microbiome of 106 infants infected with RSV. Only *H. influenzae* (*H. flu*) showed a significant association between relative abundance and clinical severity (a.) Airway gene expression in RSV infected infants was re-assessed in a multivariate model including H. flu. A circos plot displays disease ontologies associated with severity (b.) Shown are a subset of genes with expression patterns significantly associated (FDR<0.05, fold change>1.5) with clinical severity when H. flu is used as a variable in a multivariate model, and key disease ontologies significantly associated with severity (n=643 genes, FDR<0.05). Cell type enrichment analysis (c.) was used to examine genes significantly associated with severity including H. flu as compared to the same model without H. flu. This analysis showed adding H. flu to the model suggested increases in CD4+ and CD8+ T cell signatures, and decreases in myeloid cells and monocyte signatures.

Given the substantial impact of the presence of bacterial pathogens on airway gene expression in asymptomatic^35^ and symptomatic subjects (Table 2), and the singular association of *H. influenzae* with severity, we sought to identify severity-associated gene expression independent of the impact of *H. influenzae*. We performed multivariate analysis, similar to as described above, but including the relative abundance of *H. influenzae* in the model. We identified 643 genes significantly associated with GRSS independent of the presence of *H. influenzae* and visit age. Analysis of this gene set revealed many genes, pathways (Supplemental Figures 6-8) and cell types that were identified in our analysis not including *H. influenzae*. However, ontological analysis of this gene set identified a greater association with bacterial infection (Fig. 2B). Additionally, when including *H. influenzae*, we note that cell type signatures are attenuated for myeloid lineages and inflated for both CD4+ and CD8+ lymphocyte lineages (Fig. 2C). These data suggest that *H. influenzae* influences the clinical severity of RSV infection in infants by modifying the immune system and nature of the inflammatory response.

### Effects of Timing on Severity Biomarkers

As indicated above, the samples were collected at a single time during the acute phase of infection following the onset of clinical symptoms, and this timing appeared to have a major impact on gene expression (Table 2 and Supplemental Figures 9-12). We further interrogated gene expression biomarkers for this timing to better understand the nature of airway response to infection, and how they may relate to clinical disease severity. Interestingly, pathway analysis (Fig. 3A) indicated there was a strong signal of increasing EIF2 and mTOR signaling, and alterations in mitochondrial and oxidative phosphorylation activity, over time. Less surprisingly, there was also an indication of diminishing innate interferon signaling over time. The genes contributing to this signature included multiple intracellular IFN signaling molecules (MX1, IFIT1, STAT2), but not the ligands or receptors themselves. We performed exploratory analysis of these IFN pathway-related genes to better understand their expression patterns in our subjects. This revealed a surprising dichotomy between severe and non-severe subjects in expression of the IFN pathway genes identified in our analysis (Fig. 3B). An association with severity was likely not identified in our prior analysis, as significant increases in expression noted in non-severe subjects were only noted during the first 3-4 days following the onset of clinical symptoms, whereas little to no evidence for interferon signaling was noted in any subjects at later time points. qPCR validation of many of these genes supported a complex relationship between gene expression, clinical severity and the ontogeny of illness (Supplemental Table 3).

**Figure 3.**
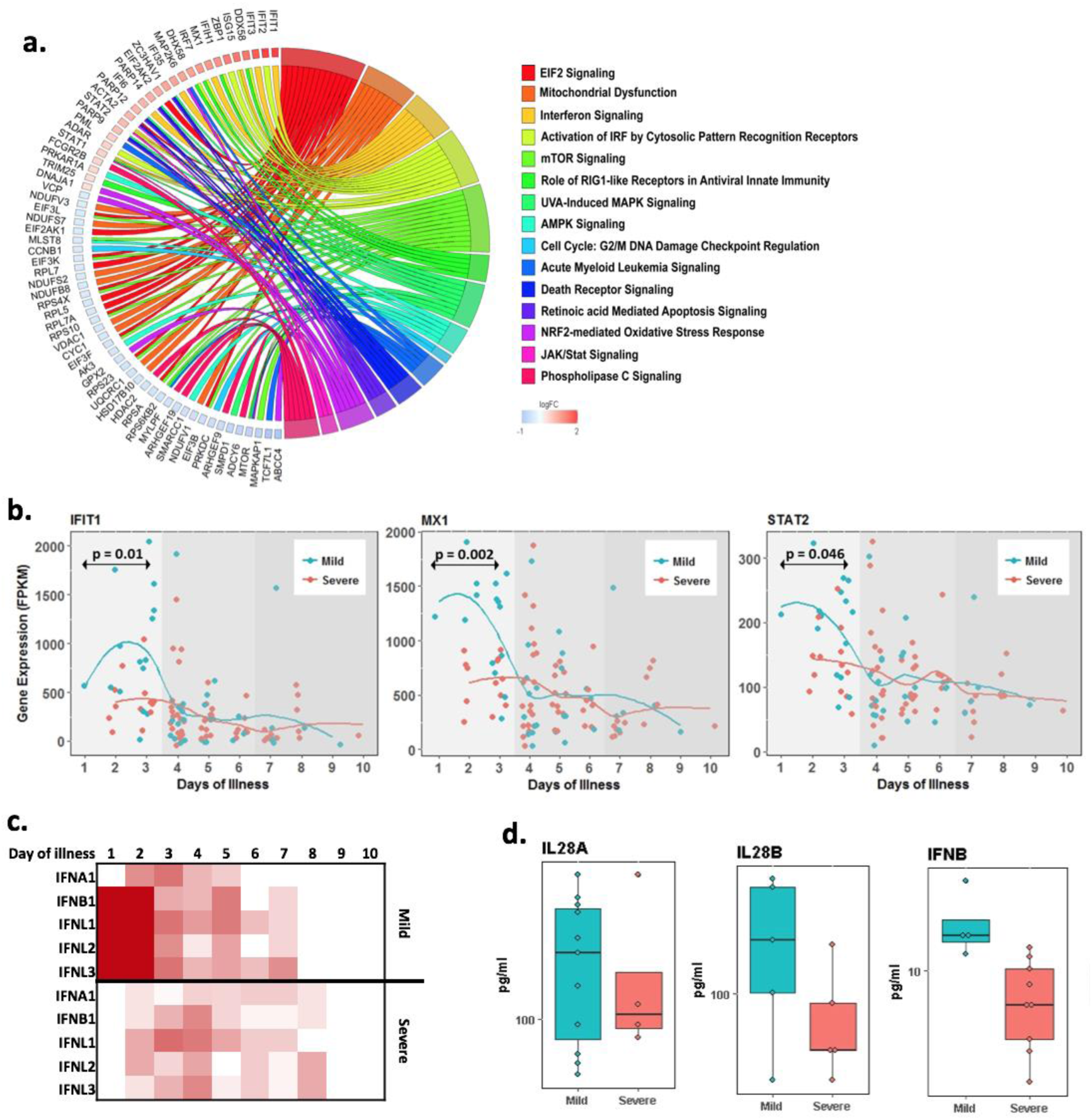
Alterations in interferon pathway activity define clinical symptoms. Airway gene expression in 106 RSV-infected infants was assessed for significant association with time since onset of clinical symptoms. A circos plot displays signaling pathway ontologies associated with severity (a.) Shown are a subset of genes with expression patterns significantly associated (FDR<0.05, fold change>1.5) with time since onset of clinical symptoms in a multivariate model, and key signaling pathway ontologies significantly associated with severity (n=692 genes, FDR<0.05). Pseudo-time series plots (b.) display gene expression patterns for key interferon signaling pathway genes significantly associated with the time since onset of clinical symptoms. IFIT1, MX1 and STAT2 display distinct patterns of temporal expression in RSV-infected infants, with significantly higher expression in infants with mild as compared to severe symptoms. Type I/III interferon ligand gene expression over time (day 1-10) was assessed separately in mild and severely affected infants (c.) Shown is the estimated expression level (more red equals higher expression) for individual ligand genes at each day following the onset of clinical symptoms. These data are consistent with an early activation in type I/III interferon pathway activity in mild subjects only. Multiplex ELISA analysis of nasal washes from RSV infected infants (d.) confirms that the production of type I/III interferon ligands is increased in infants with severe clinical symptoms.

An analysis of potential regulators for this gene expression response implicated both type 3 (IFNL1; p<10^−35^) and type 1 (IFNA2; p<10^−30^) ligands, as well as the canonical interferon-associated transcription factors IRF-3, -7 and -5 (p<10^−15^). Due to the low level of expression for many IFN ligand and receptor genes in most subjects, we removed them from our analytical data set, as part of our procedures to avoid false discovery and increase statistical power. We performed a post-hoc analysis of IFN ligand expression and found evidence for high levels of expression for both type 1 and 3 ligands (particularly IFNB1, IFNL1, IFNL2, IFNL3) in non-severe subjects, but not severe subjects (Fig. 3C). We used multiplex ELISA of nasal washings to measure airway IFN ligand production during the first few days following onset of clinical symptoms. We confirmed higher levels of IFNB and IFNL in non-severe subjects (Fig. 3D). These data suggest that interferon signaling responses and/or interferon production, within the first few days of infection, contributes to establishing clinical severity in RSV infected infants.

### Interferon Signaling Controls RSV Infection In Vitro

While interferon production is known to be a ubiquitous host response to suppress viral infection, its role in mediating the severity of RSV illness in infants is unclear, as the virus actively suppresses this host protection mechanism via the RSV NS1 and NS2 gene products^49^. Further, since our data are from the upper respiratory system, they may not accurately reflect the importance of this pathway in the lower airway as a disease mechanism contributing to clinical severity. We sought to test whether this pathway was necessary or sufficient to control the extent of RSV infection leveraging a novel in vitro model of pediatric human airway^45^. This model involves 3-D cultures of *bona fide* primary human infant/pediatric lung epithelial cells, obtained from organ donors, and differentiated at air-liquid interface^45^.

First, we demonstrated that these primary pediatric human lung epithelial (PHLE) ALI cultures are susceptible to apical infection with a laboratory strain of RSV (A2) modified to express GFP (Fig. 4A). Within 24 hours of infection, GFP+ foci are observed, and grow in number and size over 96 hours with no evidence of cell death or syncytia formation. At 48 hours post infection, significant increases in the expression of both interferons (IFNA/B/L) and interferon-stimulated genes (ISGs; MX1, IFIT1/3, RSAD2) were evident at the RNA level. We tested for the induction of expression of our severity associated biomarkers in PHLEs. Not surprisingly, as we are only modeling a portion of the in vivo responses to infection, we found that some but not all genes were regulated. We were able to demonstrate RSV infection-related increases in the expression of CCL7, IL6 and MMP19 at the RNA level.

**Figure 4.**
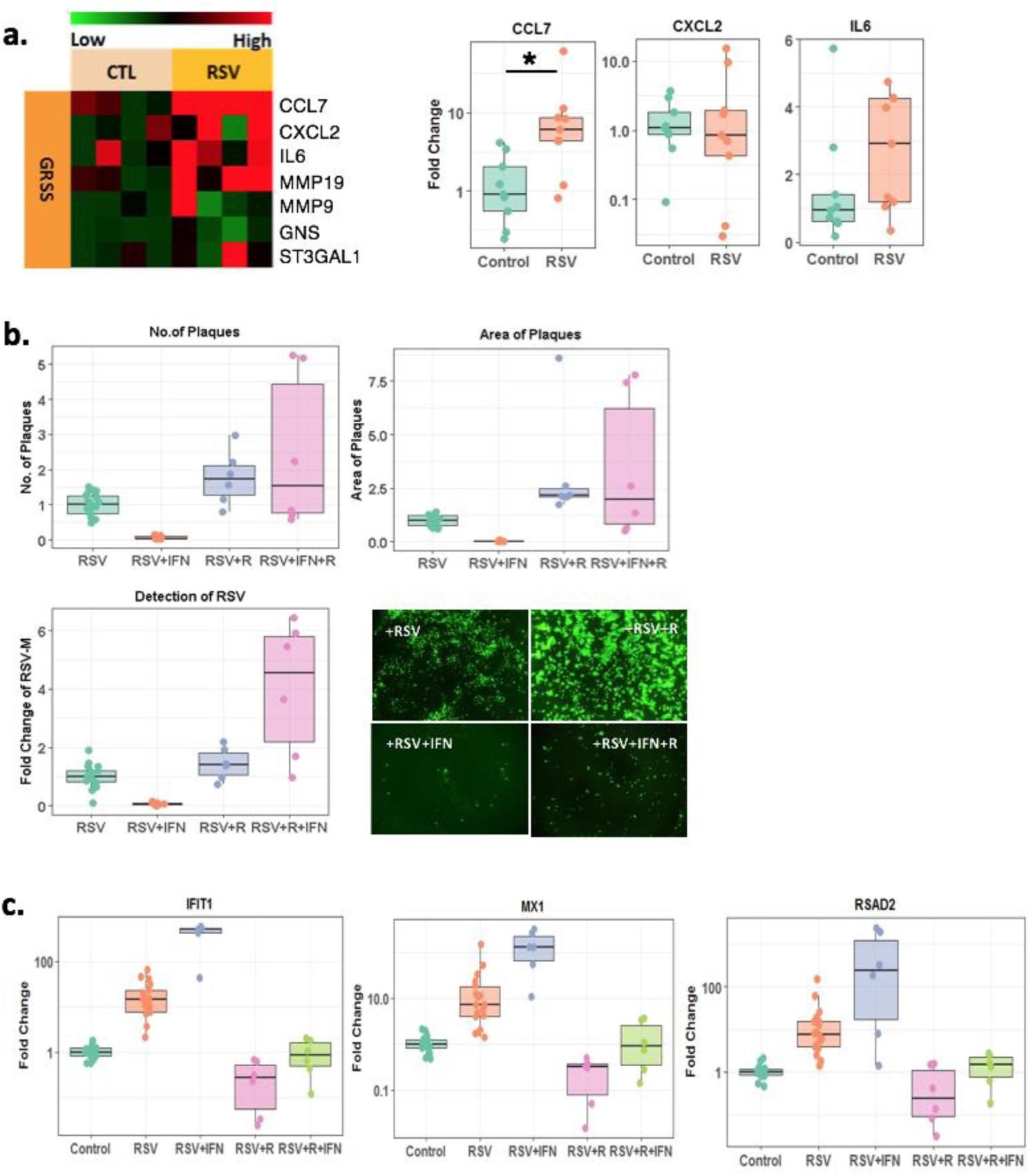
Blocking interferon signaling enhances RSV infection of primary pediatric human lung epithelial cell cultures. Infection of primary cultures of human lung epithelial (PHLE) cells differentiated at air-liquid interface with RSV emulates some gene expression responses associated with severity. (a.) Shown is a heat plot displaying relative expression for a set of genes significantly associated with the global respiratory severity score (GRSS) 48 hours after apical infection with RSV or in sham-infected controls (CTL). Also shown are box plots for multiplex ELISA for key immune mediators confirming increased production of CCL7 and IL6 following infection. (b.) Shown are the effects of RSV infection while treated with interferon ligand alone (RSV+IFN), interferon signaling inhibitor ruximitinib alone (RUX; RSV+R) or the two together (RSV+R+IFN). IFN treatment blocks the number and size of infection plaques, as well as the production of viral transcript, while blocking IFN signaling is sufficient to potentiate infection. (c.) Box plots for the expression of interferon signaling genes that are significantly associated with GRSS (IFIT1, MX1, RSAD2) shows their expression are increased by RSV infection and/or IFN treatment, but not in the presence of RUX. These data model the responses we observe in infants during acute RSV infection.

We tested the effects of gain- and loss-of-function manipulation of IFN activity during RSV infection of PHLE cells (Fig 4B). As expected, pretreatment of PHLE cells with type I (Figure 4B, C) and type III (Supplemental Figure 13-14) IFN ligand completely inhibited the extent of viral infection. We also examined the effects of blocking IFN signaling upon RSV infection in PHLE cells. The IFN receptor/STAT1 signaling blocker ruximitinib (RUX) significantly potentiated the extent of infection based upon number (2-fold, P<0.05) and size of plaques (2-fold, P<0.05), and based upon viral transcript levels (1.5-fold, P<0.05). RUX also potentiated ISG expression in RSV infected PHLE cells. Similar response were noted when cells were treated with the IFN receptor/signaling blocker TPCA (Supplemental Figure 14). Critically, but not surprisingly, RUX potentiated viral infection but did not block IFN ligand production (Fig. 4C). These data indicate IFN signaling is necessary and sufficient to suppress the expression of correlates of RSV-infected infant clinical severity in primary cultures of pediatric lung epithelial cells.

## Discussion

Our studies address a critical knowledge gap in our understanding of the mechanisms involved in infant respiratory disease, in that we rarely have had the opportunity to comprehensively interrogate resident cell responses in the target organ. We applied our recently developed methods that facilitate comprehensive molecular analysis of the airways, amenable to any infant subject, whether clinically well^35^ or during illness^47^. As applied here to infant subjects during the acute phase of RSV illness, our data supports the utility of this biospecimen as a reasonable surrogate for pathophysiological responses occurring in the lower airway (Fig. 1). In addition, they provide novel insight into disease-related responses related to the severity of clinical symptoms (Fig. 1), the presence of co-occurring microbes (Fig. 2) and the timeline of clinical symptoms (Fig. 3). We also leveraged a novel, physiological in vitro system that emulates the cellular responses to viral infection in pediatric human lung epithelial cells (PHLE). Using this model system, we validated the relationship between RSV-induced IFN-related intracellular signaling and the severity of infection, in relevant target cells for this virus (Fig. 5).

When identifying gene expression correlates of severity in this population, we were not surprised to identify many chemokines that are reflective of a more robust inflammatory response (Fig. 1). Somewhat more insightful is the ontological analysis that implicates robust changes in IL17 signaling in severe patients, when compared to those with milder forms of illness, specifically in the airway. Likewise, the data implicate airway changes in IL-1, -8, and 10 signaling, and alterations in matrix metalloproteinase activity. Of additional novel insight, our data suggests the nature of the inflammatory response in the airway differs in infants with severe RSV-associated illness. In particular, our data indicates an enrichment in CD14+ and CD133+ myeloid cells. These observations warrant further investigation to test them for potential mechanistic roles.

We were interested in understanding the role of the airway microbiota in defining clinical responses to RSV in infants. We failed to find a strong association between the presence/absence of microbial pathogens and severity. However, we confirmed a report published during our studies indicating a higher burden of *H. influenzae* in infants with severe clinical symptoms requiring hospitalization (Fig. 2). Although other taxa were associated with airway gene expression, we were somewhat surprised to find no other OTUs were significantly associated with severity. It is worth knowing, 16S ribosomal RNA sequencing was not able to identify *S. pneumoniae* at the species level. Given our prior studies revealing a large impact of the microbiota upon gene expression in the airway, we studied the inter-relationships between the airway microbiome, airway gene expression and clinical severity. In general, these analyses confirmed that the existence of known pathogens, particularly bacteria, are strongly associated with gene expression in asymptomatic^35,47^ and symptomatic^47^ infants. We considered the specific effect of the microbiota/H. flu on the relationship between gene expression and RSV severity. The results are consistent with H. flu altering the nature of the inflammatory response, from myeloid-predominant to CD4/8 T lymphocyte-predominant. These data provide a currently untested hypothesis for a potential mechanism whereby *H. influenzae* contributes to severe responses at the molecular and cellular level, which is a focus of current investigation.

Our identification of distinct, time-dependent changes in IFN signaling when comparing severely affected versus non-severely affected infants is both novel and consistent with the fundamental biology of viral responses. Perhaps this was not readily apparent in prior studies focusing on severely affected infants, as compared to non-infected infants, as they both show limited evidence for activation of this pathway. In this way, our studies, focused upon clinical severity in a cohort where all were infected with RSV, provides opportunity for different insights. We also draw attention to the narrow time window in which differences in IFN between these groups is evident (Fig. 4). Even in our own data, differences in expression do not reach the level of statistical significance without restricting our analyses to the first few days following the onset of clinical symptoms.

In vitro, our data indicating that IFN is both necessary and sufficient to control RSV infection of mammalian cells is not particularly novel, but validates the hypothesis generated by our in vivo data. However, we feel that it is important to demonstrate this mechanism is causal for infection in a truly physiological model of *bona fide* human pediatric lung epithelial cells. These in vitro data also partially reveal the complexity of the in vivo response, in that only a subset of gene expression correlates of severity are validated. We suggest other severity-associated gene expression responses require, or result from, additional cell types (e.g., leukocytes), and likely to be modified by the microbial community. Modeling these states and responses are one focus of future experimentation.

We should emphasize that our primary observation is not that severity is associated with IFN ligand production, but with estimated levels of signaling activation in the airway. This distinction could be due to low levels of ligand production, even in infants with mild clinical symptoms. However, we could show differences in IFN ligand expression at both the RNA and protein levels (Fig. 1), after our initial discovery phase of RNAseq data analysis. Conversely, the distinction could support the logical conclusion that downstream signaling is critical to the protective effects of viral-induced IFN production. This interpretation is supported by our in vitro data (Fig. 5), where blocking IFN receptor signaling is sufficient to potentiate widespread viral infection, even in the presence of IFN ligand expression. Regardless, induction of type I and type III IFN does occur in vivo and in vitro. The relative role of type I versus type III IFN in vivo is not clear at this time. Both type I and type III IFN appear to be expressed in non-severe subjects during the acute phase of illness (Fig. 4) and can be induced by viral infection in vitro.

Our evaluation of human biospecimen-derived RNAseq data has utilized multiple analytical and statistical approaches, in an effort to thoroughly interrogate the in vivo responses they reflect. Critically, we have applied appropriately conservative corrections for multiple testing in each case. We believe that use of each of these analytical approaches is justified, as they each provide distinct yet important insight. Univariate analyses are used to estimate the direct associations between one covariate and individual genes. These analyses are easy to perform and interpret and do not suffer from potential collinearity issues among covariates. As such, they are useful to initially explore the overall pattern of associations between clinical covariates and transcriptome profiles. On the other hand, a multivariate model considers the influence of several covariates on gene expression *simultaneously*; therefore, the estimated association between the response variable and a given covariate is less likely to be affected by unmodeled interdependences. Arguably, a multivariate model is more natural than multiple univariate models because it more closely approximates the complex biological processes that collectively influence the transcriptome. That being said, more caution must be exercised when using complex multivariate models. When uninformative covariates are included in a multivariate model, they may mask true associations and reduce the statistical power to detect informative associations. A more serious issue is the potential collinearity among the covariates, which may prevent modeling certain combinations of covariates and produce unstable estimates. Therefore, we believe it is a practical strategy to first use univariate analyses to identify a subset of important covariates, and then use them to build a lean and robust multivariate model.

We acknowledge this study is not without limitations. We have focused our efforts in identifying gene expression responses that distinguish infants severely affected by RSV infection as compared to infants who are not severely ill and have not included non-infected controls. Therefore, we are unable to address the specificity of the responses we observe and report. Other studies by our group (Storch and unpublished) suggest that responses to any viral infection can be identified but occur at a much smaller magnitude than those described here. Another limitation is the associative nature of most of the data presented. One must use caution and resist presuming that the in vivo correlative observations we report are mechanistic, rather than hypothesis generating. In some cases, we were able to show a mechanistic relationship between specific genes/pathway and the magnitude of viral infection (Fig. 5). Importantly, this in vitro model provides a physiologically relevant system to further pursue mechanistic hypotheses resulting from analysis of our in vivo data. However, this model also has limitations in that it currently isolates epithelial cell responses, while our data clearly suggests involvement of leukocytes and monocytes in vivo.

## Acknowledgements

Complete molecular and microbiota data for these studies is available in dbGaP (phs001201.v2.p1). The authors would like to thank the Aspires team for critical assistance with subject recruitment and sample collection. This project has been funded with Federal funds from the National Institute of Allergy and Infectious Diseases, National Institutes of Health, Department of Health and Human Services, under Contract No. HHSN272201200005C. Finally, we are indebted to the patients and families who agreed to participate in these studies.

