## Supplemental document for "Insufficiency in airway interferon activation defines clinical severity to infant RSV infection"

**Data Supplement for:**

Supplemental Tables: 3

Supplemental Figures: 14

**Supplemental Table 1.** qPCR was used to validate the expression patterns associated with GRSS and correlated to RNA-seq. Shown are p-values of rank correlation test of association based upon GRSS and qPCR data, comparing gene expression determined by qPCR to RNA-seq.

| Gene name | **p-value of correlation of GRSS** | **p-value of correlation of RNA-seq** |
|---|---|---|
| CCL7 | **0.0494** | **0.0120** |
| CD3E | 0.1258 | **0.0009** |
| IL6 | **0.0081** | **1.62E-06** |
| GNS | 0.7028 | **0.0133** |
| VCAN | **0.0028** | **6.58E-08** |

**Supplemental Table 2.** Significant OTUs from univariate analysis for microbiome data

| OTU | GRSS p-value unadjusted | GRSS p-value BH adjusted |
|---|---|---|
| **g__Haemophilus,s__influenzae** | **0.00001** | **0.00178** |
| g__Haemophilus,s__parainfluenzae | 0.00161 | 0.11893 |
| g__Actinomyces,s__ | 0.00496 | 0.18360 |
| g__Chryseobacterium,s__ | 0.00440 | 0.18360 |
| g__Atopobium,s__ | 0.01920 | 0.29817 |
| g__Granulicatella,s__ | 0.02015 | 0.29817 |
| g__Lactococcus,s__ | 0.01751 | 0.29817 |
| g__Acinetobacter,Unknown | 0.01679 | 0.29817 |
| g__Haemophilus,Unknown | 0.01398 | 0.29817 |
| g__Klebsiella,Unknown | 0.01224 | 0.29817 |
| g__Dietzia,s__ | 0.04531 | 0.50277 |
| g__Ochrobactrum,s__ | 0.03769 | 0.50277 |
| g__Diaphorobacter,s__ | 0.04671 | 0.50277 |
| g__Neisseria,s__subflava | 0.04756 | 0.50277 |
| | **Visit Age p-value unadjusted** | **Visit Age p-value BH adjusted** |
| **g__Porphyromonas,s__** | **0.00000** | **0.00017** |
| **g__Staphylococcus,Unknown** | **0.00000** | **0.00017** |
| **g__Neisseria,s__subflava** | **0.00001** | **0.00039** |
| **g__Staphylococcus,s__epidermidis** | **0.00001** | **0.00040** |
| **g__Granulicatella,s__** | **0.00008** | **0.00235** |
| **g__Prevotella,s__nanceiensis** | **0.00092** | **0.02276** |
| **g__Actinomyces,s__** | **0.00263** | **0.04354** |
| **g__Rothia,s__dentocariosa** | **0.00250** | **0.04354** |
| **g__Neisseria,Unknown** | **0.00265** | **0.04354** |
| g__[Prevotella],s__ | 0.00468 | 0.06298 |
| g__Veillonella,Unknown | 0.00428 | 0.06298 |
| g__Lactococcus,s__ | 0.00993 | 0.11304 |
| g__Haemophilus,Unknown | 0.00988 | 0.11304 |
| g__Luteimonas,s__ | 0.01129 | 0.11937 |
| g__Veillonella,s__parvula | 0.01692 | 0.14726 |
| g__Leptotrichia,s__ | 0.01677 | 0.14726 |
| g__Ochrobactrum,s__ | 0.01516 | 0.14726 |
| g__Capnocytophaga,s__ | 0.01892 | 0.14922 |
| g__Parvimonas,s__ | 0.01916 | 0.14922 |
| g__Sphingobium,s__ | 0.02066 | 0.15290 |

| | | |
|---|---|---|
| g__Neisseria,s__cinerea | 0.02297 | 0.16189 |
| g__Prevotella,s__ | 0.03000 | 0.20185 |
| g__Pseudomonas,s__fragi | 0.03164 | 0.20358 |
| g__Achromobacter,s__ | 0.03320 | 0.20471 |
| g__Campylobacter,s__ | 0.04679 | 0.27701 |
| | **Days Since Onset p-value unadjusted** | **Days Since Onset p-value BH adjusted** |
| g__Microbacterium,Other | 0.02139882 | 0.6332523 |
| g__Rothia,s__dentocariosa | 0.02567239 | 0.6332523 |
| g__Prevotella,s__melaninogenica | 0.01167967 | 0.5205303 |
| g__Streptococcus,s__agalactiae | 0.01406839 | 0.5205303 |
| g__Anaerococcus,s__ | 0.01040704 | 0.5205303 |
| g__Pandoraea,s__ | 0.00708746 | 0.5205303 |
| | **Mode of Delivery p-value unadjusted** | **Mode of Delivery p-value BH adjusted** |
| g__Staphylococcus,s__epidermidis | 0.04344 | 0.88539 |
| g__Bulleidia,s__moorei | 0.00385 | 0.28475 |
| g__Leptotrichia,s__ | 0.01122 | 0.44680 |
| g__Gluconacetobacter,s__ | 0.01208 | 0.44680 |
| g__Haemophilus,s__parainfluenzae | 0.04228 | 0.88539 |
| g__Meiothermus,s__ | 0.04106 | 0.88539 |
| g__Klebsiella,Unknown | 0.00227 | 0.28475 |
| | **Antibiotic Use p-value unadjusted** | **Antibiotic Use p-value BH adjusted** |
| g__Corynebacterium,s__kroppenstedtii | 0.02385 | 0.50421 |
| g__Dietzia,s__ | 0.04701 | 0.55667 |
| g__Propionibacterium,s__granulosum | 0.04576 | 0.55667 |
| g__[Prevotella],s__ | 0.03108 | 0.55505 |
| g__Capnocytophaga,s__ | 0.03375 | 0.55505 |
| g__Streptococcus,s__infantis | 0.01908 | 0.50421 |
| g__Peptostreptococcus,s__ | 0.00697 | 0.45719 |
| g__Brevundimonas,s__diminuta | 0.02215 | 0.50421 |
| g__Enhydrobacter,s__ | 0.04966 | 0.55667 |
| g__Pseudomonas,s__viridiflava | 0.00927 | 0.45719 |
| g__Meiothermus,s__ | 0.00044 | 0.06524 |
| g__Thermus,s__ | 0.01714 | 0.50421 |

**Supplemental Table 3.** qPCR was used to validate the expression patterns associated with onset of illness and correlated to RNA-seq. Shown are p-values of rank correlation test associated with onset of illness and qPCR data, comparing gene expression determined by qPCR to RNA-seq.

| **Gene name** | **p-value of correlation of onset of illness** | **p-value of correlation of GRSS** | **p-value of correlation of GRSS (day 0-3)** |
|---|---|---|---|
| **IFIT1** | **0.0037** | 0.3276 | **0.0061** |
| **HERC5** | **0.0063** | 0.5526 | **0.0325** |
| **RSAD2** | **0.0214** | 0.5575 | **0.0464** |
| **IFNL1** | **0.0160** | 0.8143 | **0.0464** |
| **IFNL2** | **0.0055** | **0.0264** | 0.0977 |

### Supplemental Figure Legends

**Supplemental Figure 1. Sequencing quality assessment.** Details are provided for total number of sequencing reads generated (a), genome mapping rate (b), and the proportion of the genome for which transcripts were detected (c), for each of the samples.

**Supplemental Figure 2. Candidate epithelial and inflammatory cell marker expression.** Individual genes are indicated as rows, and individual subjects as columns. Relatively high expression is indicated by red and low expression is indicated by green. Expression estimates of candidate epithelial cell and leukocyte genes are presented. The data indicate consistently high levels of expression for epithelial cell markers including prototypical nasal epithelial cell genes (BPIFs), general epithelial cell genes (CDH1) and some mucosal epithelial cell markers (MUC1/4/16/20).

**Supplemental Figure 3. Gene expression associated with disease severity in RSV-infected infants.** Shown are normalized expression levels for the 252 genes associated with disease severity. Individual genes are indicated as columns, and individual subjects are indicated as rows. Relatively high expression is indicated by red and low expression is indicated by green. Samples are grouped by global severity score (GRSS).

**Supplemental Figure 4. Regulators associated with clinical severity in RSV-infected infants.** Upstream regulators defined by the set of 252 genes with expression associated with clinical severity were identified using Ingenuity Pathway Analysis (IPA). Shown are the 25 upstream regulators with the lowest significant p-values. Orange bars indicate pathways predicted to be activated; blue bars indicate pathways predicted to be inhibited; grey bars indicate pathways without a predicted directionality. Target genes for each regulator are listed.

**Supplemental Figure 5. Defining *Haemophilus influenzae* burden.** Pairwise scatter plot of abundance from 16s rRNA sequencing shows that *Haemophilus influenzae* and *Haemophilus* unknown species are highly correlated (a.). Detection of H. *influenzae* using TAC in 106 infants infected with RSV is shown in (b), detection of *Haemophilus* unknown species in 106 infants infected with RSV is shown in (c).

**Supplemental Figure 6. Biological pathways associated with *Haemophilus* in RSV-infected infants.** IPA pathway was used to identify biological functions represented the set of

643 genes with expression associated with *Haemophilus*. Shown are a subset of genes with expression patterns significantly associated (FDR<0.05, fold change>1.5) with *Haemophilus*, following adjustment for other variables in a model.

**Supplemental Figure 7. Biological functions associated with *Haemophilus* in RSV-infected infants.** IPA pathway was used to identify biological functions represented the set of 643 genes with expression associated with *Haemophilus*. Shown are a subset of genes with expression patterns significantly associated (FDR<0.05, fold change>1.5) with *Haemophilus*, following adjustment for other variables in a model.

**Supplemental Figure 8. Regulators associated with *Haemophilus* in RSV-infected infants.** Upstream regulators defined by the set of 643 genes with expression associated *Haemophilus* were identified using IPA. Shown are the 25 upstream regulators with the lowest significant p-values. Orange bars indicate pathways predicted to be activated at later times; blue bars indicate pathways predicted to be inhibited at later times; grey bars indicate pathways without a predicted directionality. Target genes for each regulator are listed.

**Supplemental Figure 9. Gene expression associated with time since onset of clinical symptoms in RSV-infected infants.** Shown are normalized expression levels for the 728 genes associated with time since onset of clinical symptoms. Individual genes are indicated as columns, and individual subjects as rows. Relatively high expression is indicated by red and low expression is indicated by green. Samples are grouped by days since onset of illness.

**Supplemental Figure 10. Disease ontologies associated with time since onset of clinical symptoms in RSV-infected infants.** IPA pathway was used to identify diseases represented the set of 728 genes with expression associated with day of illness. Relationships between pathway and individual differentially expressed genes are shown. Shown are a subset of genes with expression patterns significantly associated (FDR<0.05, fold change>1.5) with time since onset of clinical symptoms, following adjustment for other variables in a multivariate model.

**Supplemental Figure 11. Biological functions associated with time since onset of clinical symptoms RSV-infected infants.** IPA pathway was used to identify pathways represented the set of 728 genes with expression associated with define clinical symptoms. Relationships between pathway and individual differentially expressed genes are shown. Shown are a subset of genes with expression patterns significantly associated (FDR<0.05, fold change>1.5) with time since onset of clinical symptoms, following adjustment for other variables in a multivariate model.

**Supplemental Figure 12. Regulators associated with time since onset of clinical symptoms in RSV-infected infants.** Upstream regulators defined by the set of 728 genes with expression associated with day of illness were identified using IPA. Shown are the 25 upstream regulators with the lowest significant p-values. Orange bars indicate pathways predicted to be activated at later times; blue bars indicate pathways predicted to be inhibited at later times; grey bars indicate pathways without a predicted directionality. Target genes for each regulator are listed.

**Supplemental Figure 13. Effect of IFN signaling inhibitor TCPA on RSV infection of primary pediatric human lung epithelial cell cultures.** ***Top,*** Shown are fluorescent images of ALI PHLE cultures infected with RSV-GFP in the absence (left) or presence (right) of the IFN signaling inhibitor TPC. ***Bottom,*** Quantitative assessment of the effects of TCPA on RSV infection in PHLE, in the absences of presence of TCPA,  as defined by the number of GFP+ plaques, the total area of GFP+ plaques and the level of RSV-M transcript.

**Supplemental Figure 14. The effects of type III IFN on RSV infection of primary pediatric human lung epithelial cell cultures.** (a) Shown are the effects of RSV infection while treated with type III interferon ligand alone (RSV+IL28), interferon signaling inhibitor ruximitinib alone (RUX; RSV+R) or the two together (RSV+R+IL28). IFN treatment blocks the number and size of infection plaques, as well as the production of viral transcript, while blocking IFN signaling is sufficient to potentiate infection. (b) Box plots for the expression of interferon signaling genes that are significantly associated with GRSS (IFIT1, MX1, RSAD2) shows their expression are increased by RSV infection and/or IFN treatment, but not in the presence of RUX.

### Supplemental Figure 1

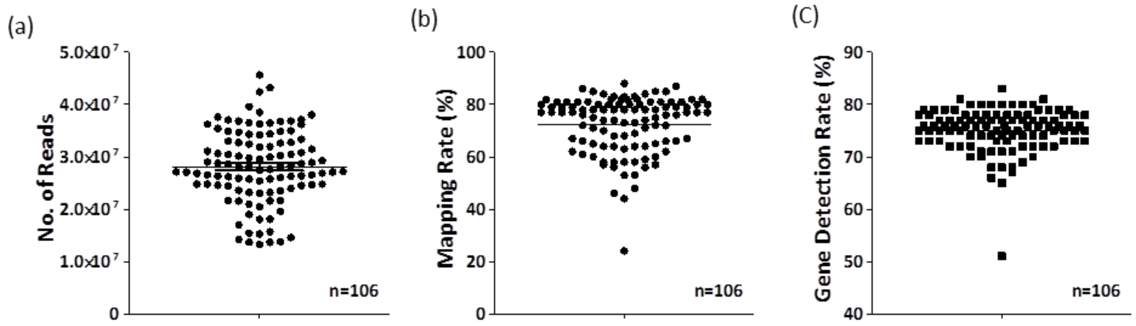

**Supplement Figure 2.**

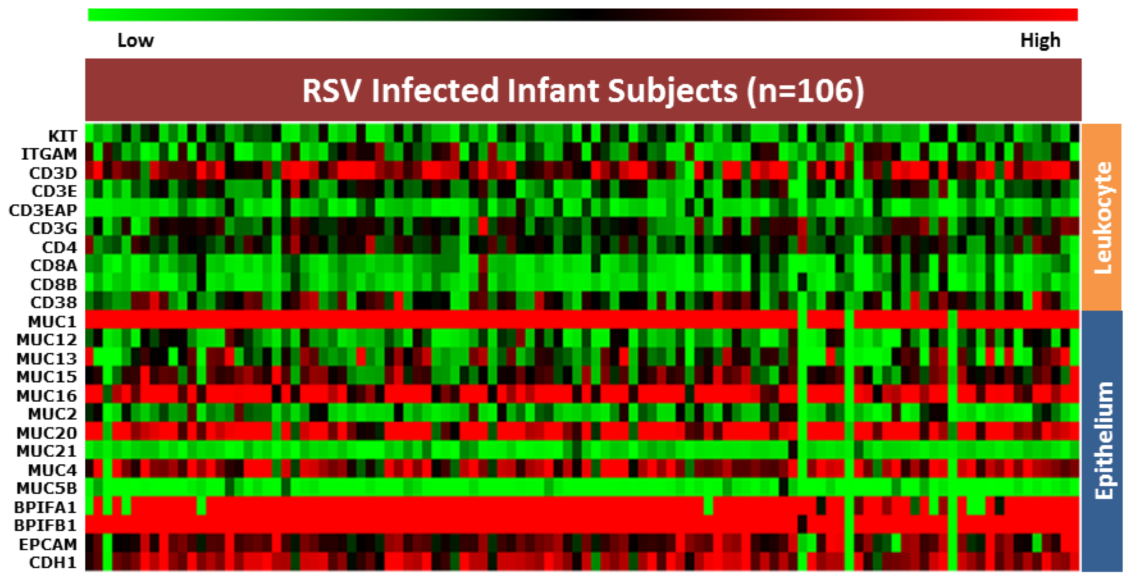

**Supplemental Figure 3.**

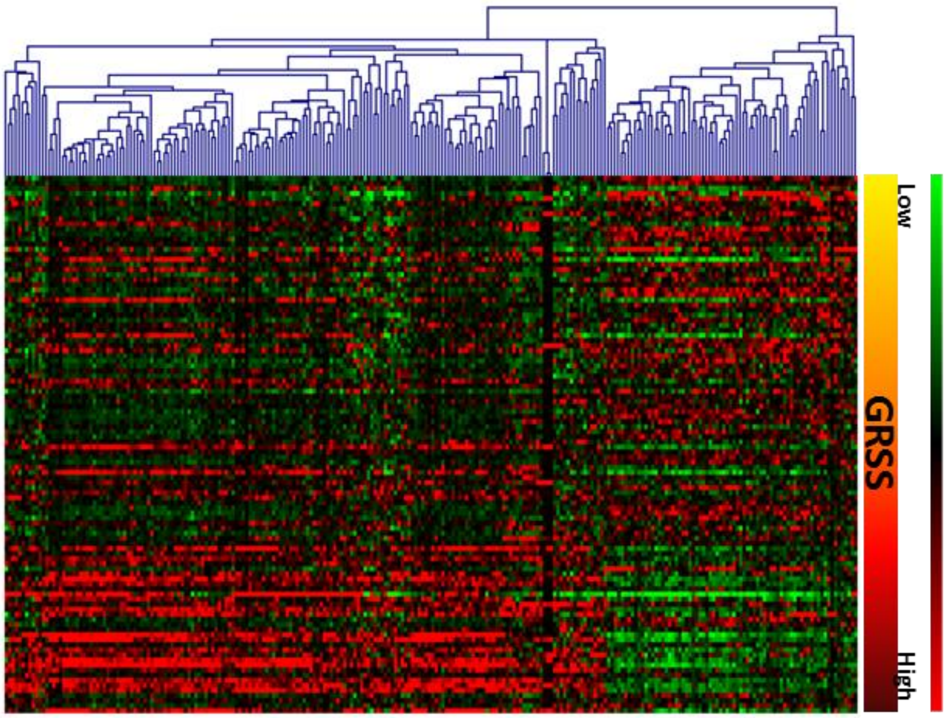

### Supplemental Figure 4.

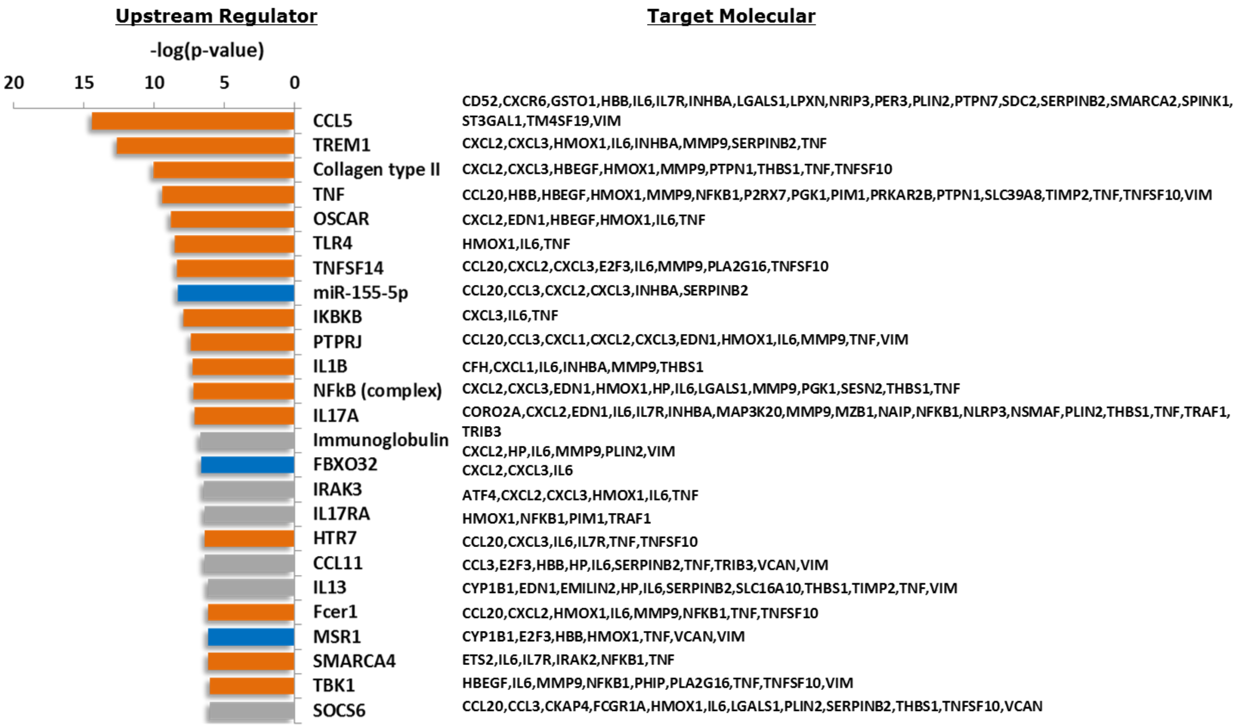

**Supplemental Figure 5.**

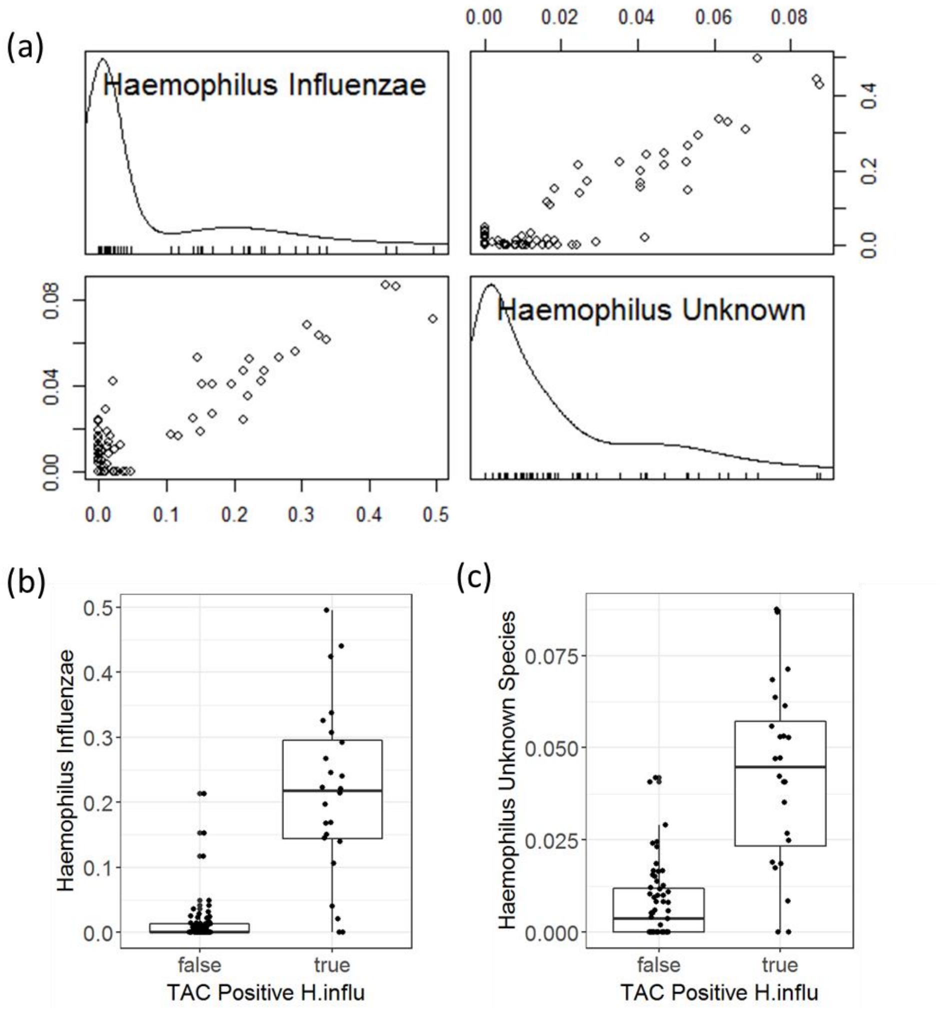

**Supplemental Figure 6.**

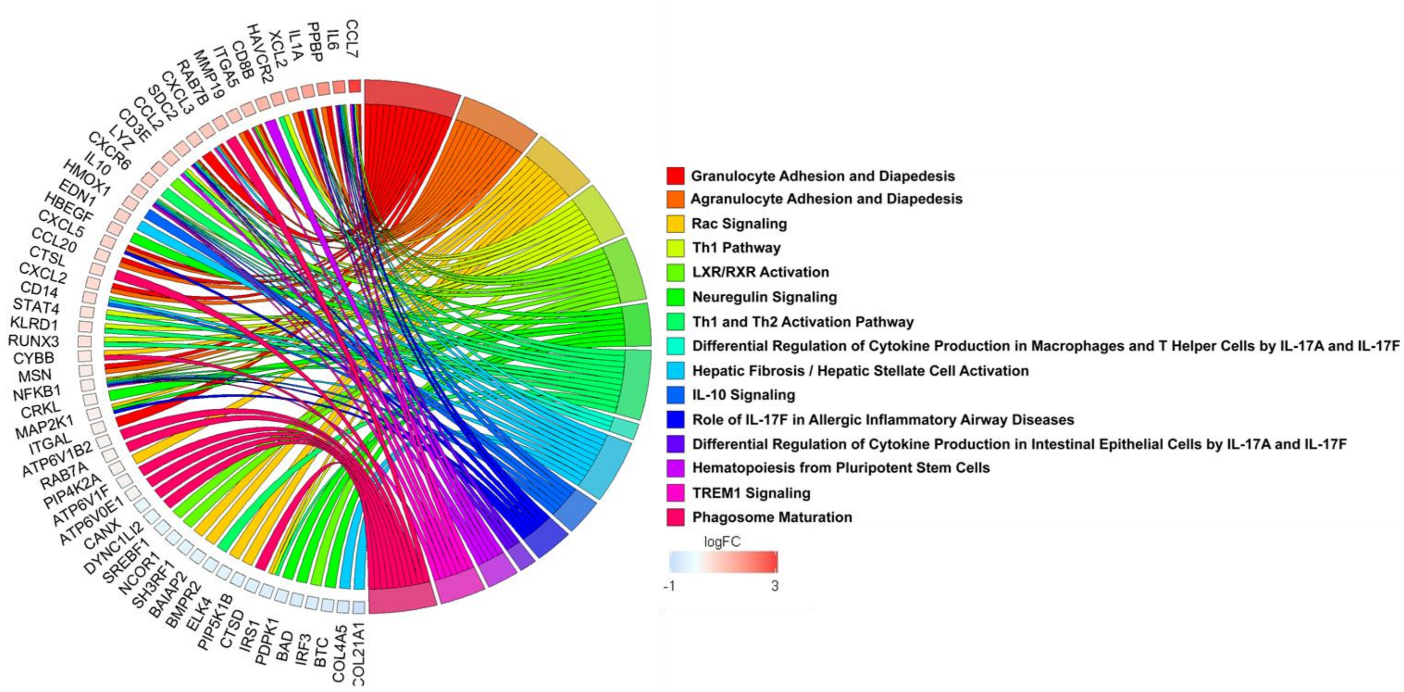

**Supplemental Figure 7.**

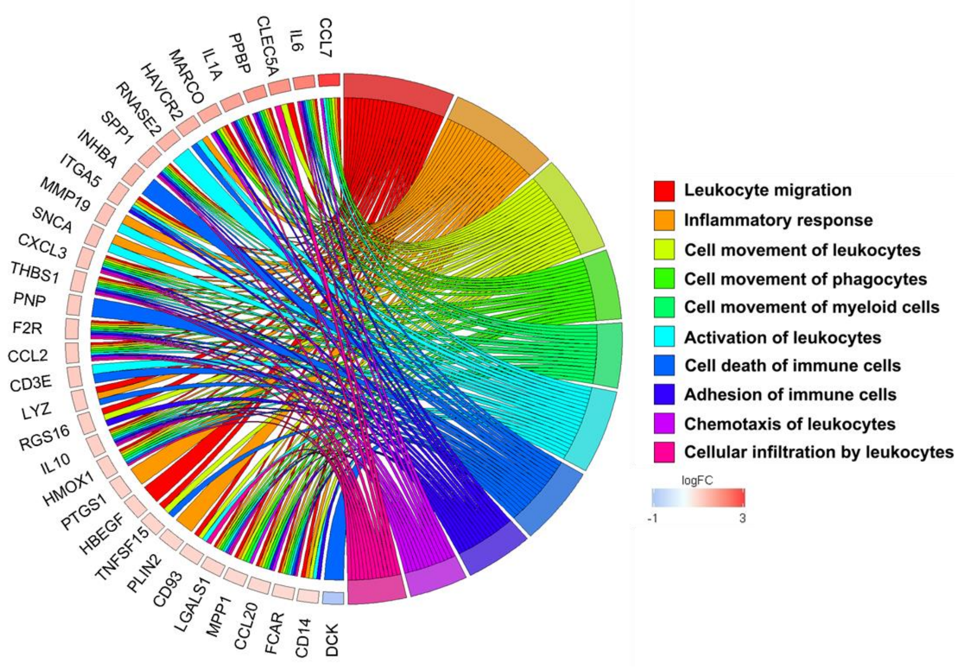

### Supplemental Figure 8.

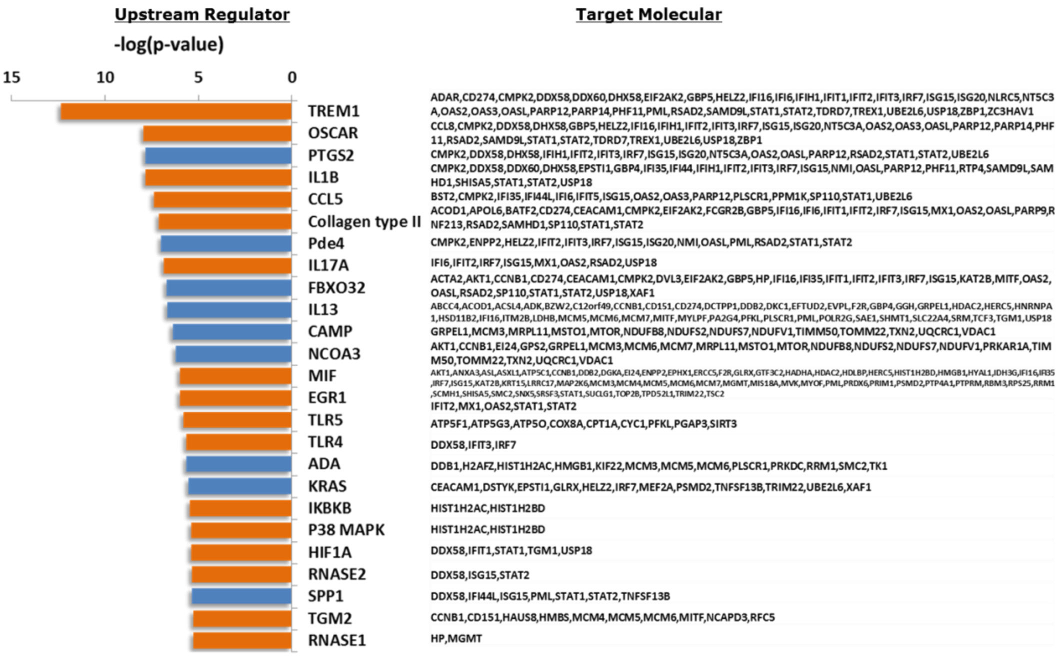

**Supplemental Figure 9.**

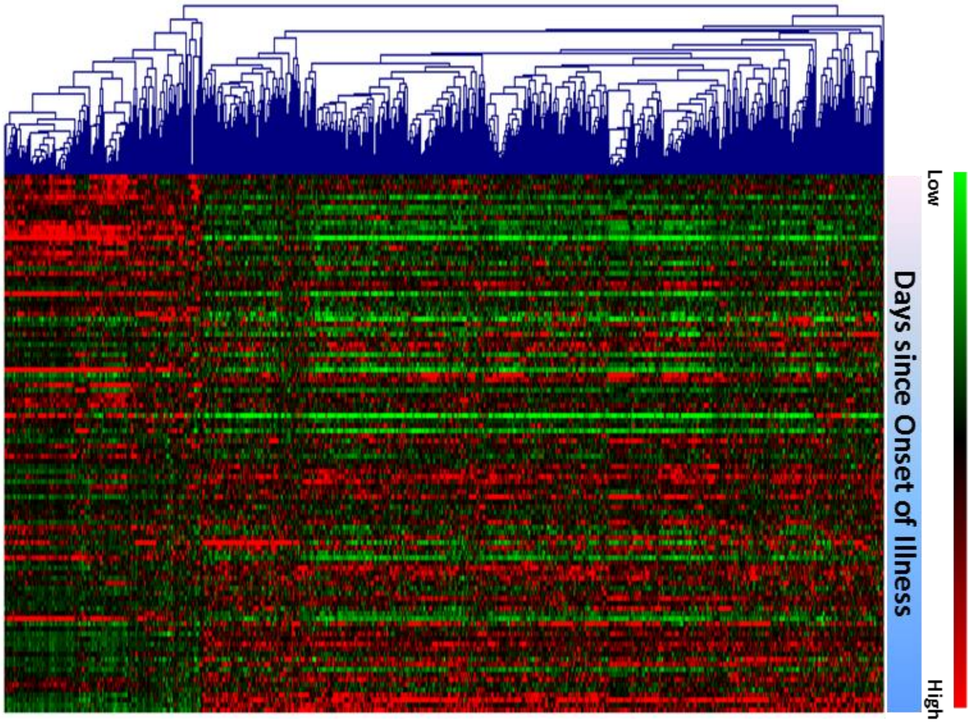

**Supplemental Figure 10.**

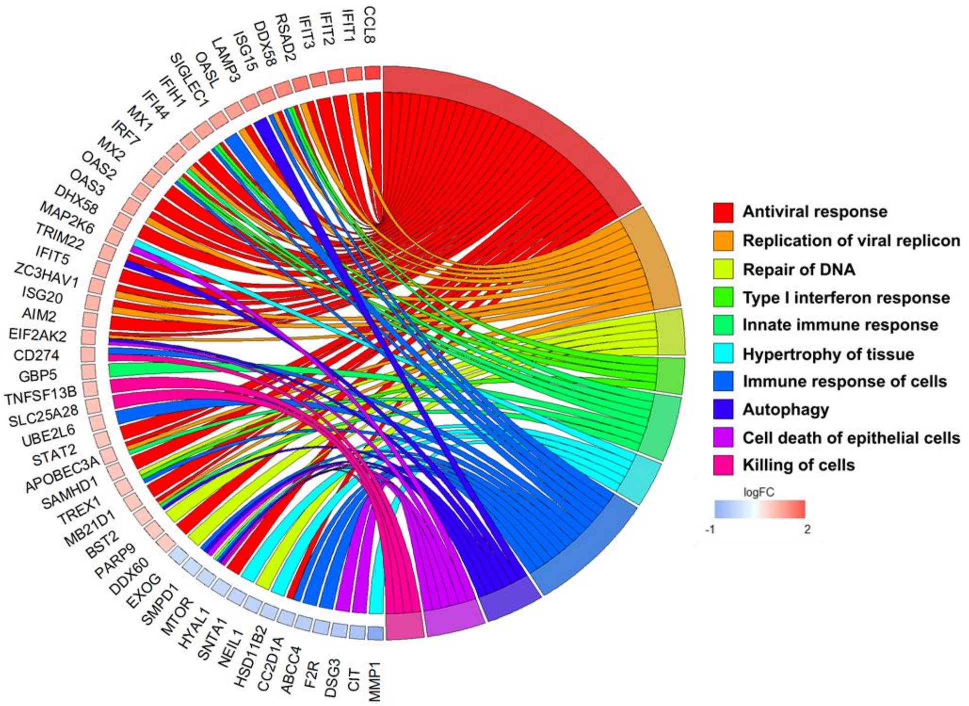

**Supplemental Figure 11.**

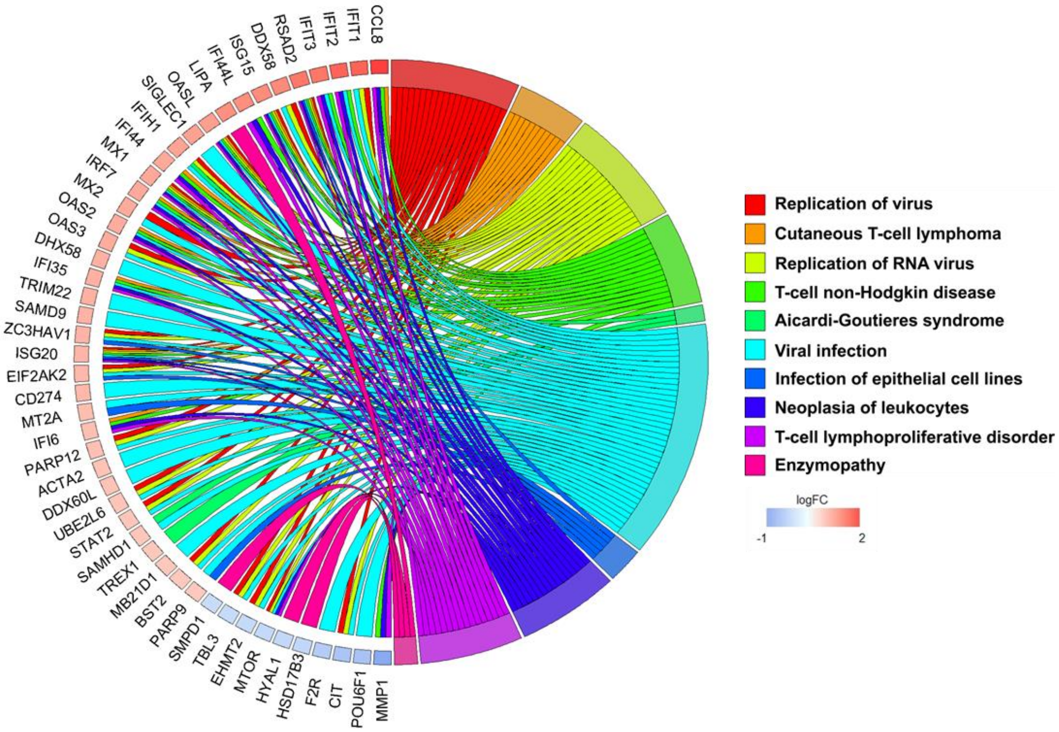

**Supplemental Figure 12.**

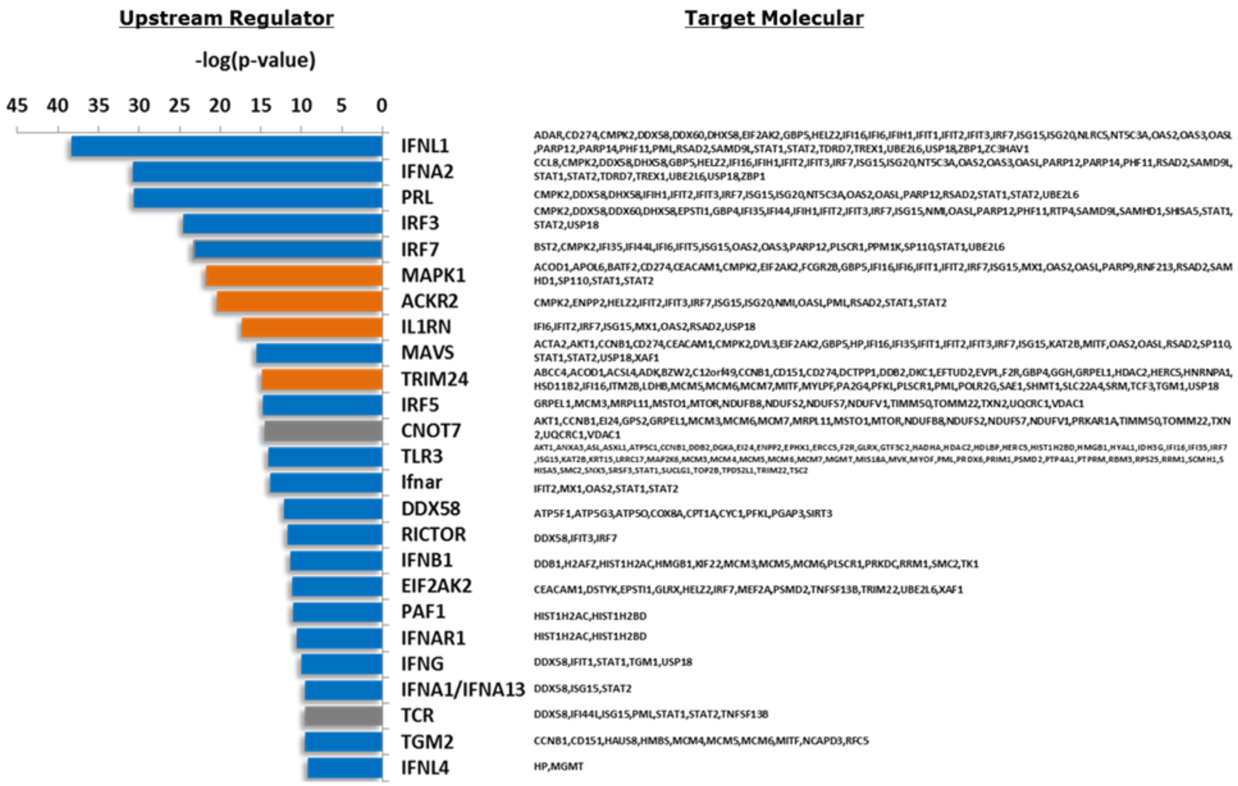

**Supplemental Figure 13.**

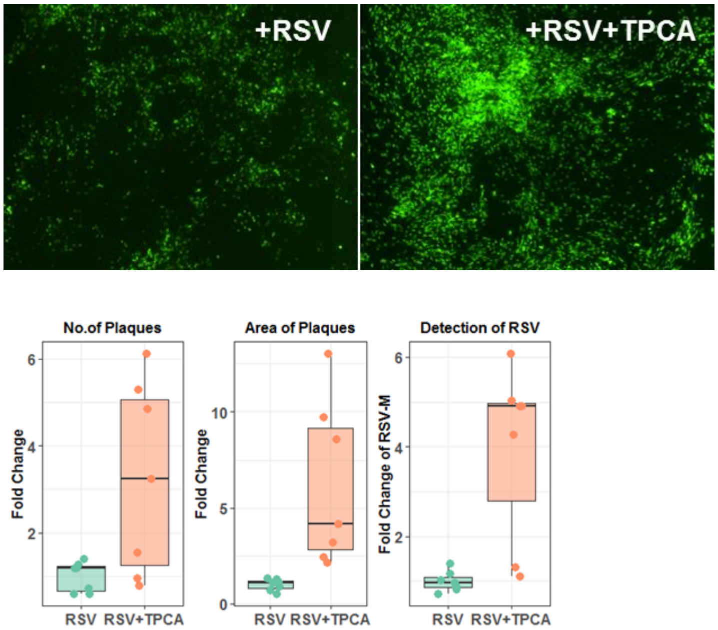

**Supplemental Figure 14.**

**a**

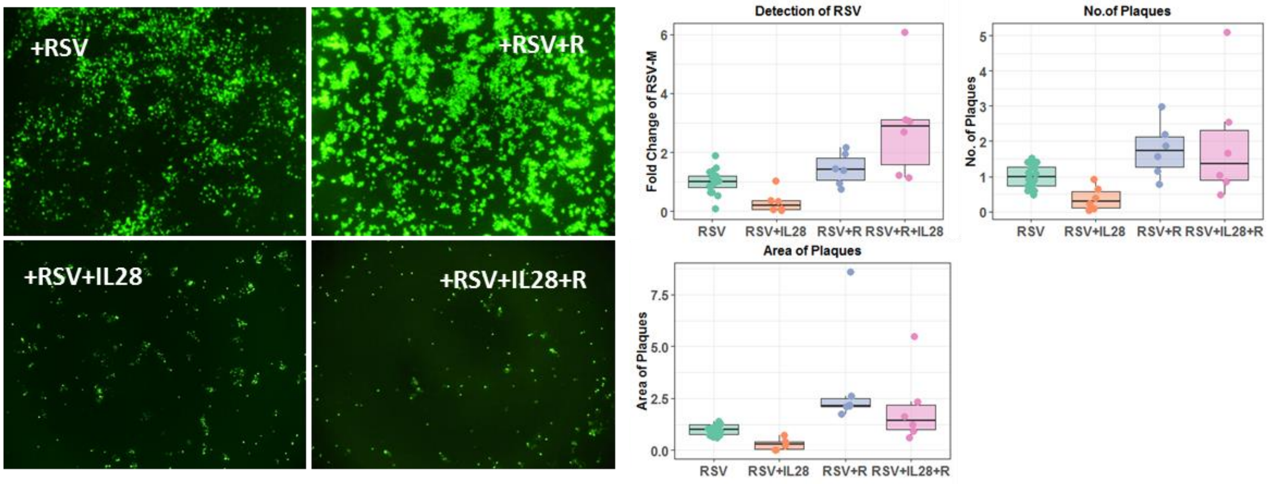


**b**

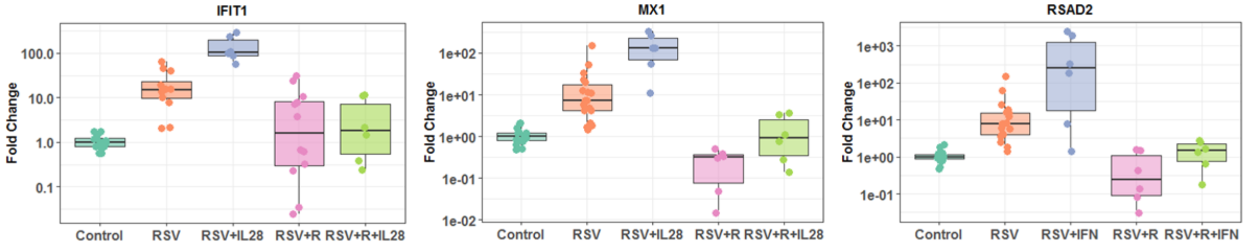
